# Facing pain is effortful: key role of the supplementary motor area and anterior midcingulate cortex

**DOI:** 10.64898/2026.04.17.719211

**Authors:** Ilaria Monti, Marie-Eve Picard, Thomas Mangin, Maxime Bergevin, Mathieu Gruet, Stéphane Baudry, A. Ross Otto, Jen-I Chen, Mathieu Roy, Pierre Rainville, Benjamin Pageaux

**Affiliations:** Centre de Recherche de L’Institut Universitaire de Gériatrie de Montréal (CRIUGM), H3W 1W4 Montréal, Québec, Canada; École de Kinésiologie et des Sciences de l’Activité Physique (EKSAP), Faculté de Médecine, Université de Montréal, H3T 1J4 Montréal, Québec, Canada; Département de psychologie, Faculté des arts et des sciences, Université de Montréal, H2V 2S9 Montréal, Québec, Canada; J-AP2S Laboratory, University of Toulon, 83041 Toulon, France; Laboratory of Applied Biology and Neurophysiology, ULB Neuroscience Institute, Université libre de Bruxelles, 1070 Brussels, Belgium; Department of Psychology, McGill University, H3A 1G1 Montreal, Québec, Canada; Alan Edwards Centre for Research on Pain, McGill University, H3A 2B4 Montreal, Québec, Canada; Département de stomatologie, Université de Montréal, H3C 3J7 Montréal, Québec, Canada; Centre interdisciplinaire de recherche sur le cerveau (CIRCA), H3T 1P1 Montréal, Québec, Canada

**Keywords:** Neuroimaging, experimental pain, visuomotor performance, perception of effort

## Abstract

Pain captures attention and interferes with executive and motor processes. In the presence of pain, increasing effort may represent a compensatory mechanism to counteract pain-related disruption and maintain task performance. In this preregistered fMRI study, we investigated neural mechanisms underlying preserved task performance during pain and increased perceived effort.

Forty right-handed participants performed a visuomotor force-matching task consisting of isometric handgrip contractions at a low and high force levels under individually calibrated painful or non-painful thermal stimulation. Thermal stimulation was applied to the left forearm, and participants rated the intensity of perceived effort after each isometric contraction.

Maintaining task performance under pain was associated with increased perceived effort and recruited brain regions involved in pain modulation and cognitive control. Region-of-interest analysis showed perceived effort was consistently linked to decreased anterior midcingulate cortex activity, whereas supplementary motor area contributions varied depending on its role in motor execution or pain processing. Across experimental condition, motor, pain-modulatory and cognitive-control regions were associated with effort perception. Independently of condition, effort perception was modulated by ventromedial prefrontal cortex and ventral striatum. These findings indicate that effort perception is a complex phenomenon reflecting brain activity within areas involved in motor, executive and valuation processes.

**Significance Statement:** This study advances our understanding of the neural mechanisms underlying task performance under pain and increased effort perception. Brain activity was measured during a visuomotor force-matching task performed in the presence or absence of pain. By contrasting task-related activity between painful and non-painful conditions, we identified regions associated with cognitive control and pain modulation involved in preserving task performance under pain. By correlating activity in regions of interest with ratings of perceived effort, we demonstrated the involvement of the supplementary motor area and midcingulate cortex in effort perception. These findings suggest that additional neural resources are recruited to maintain performance during pain and that the supplementary motor area and midcingulate cortex contribute to heightening the effort experienced.

## Introduction

Pain captures attention and disrupts executive and motor processes, impairing task performance during cognitive and motor tasks (e.g., Moore et al., 2012; Salomoni & Graven-Nielsen, 2012). However, task performance can sometimes be preserved despite pain interference (Smith et al., 2006; Tabry et al., 2020). We and others proposed that maintaining task performance under pain requires actively increasing effort to counteract pain-related interference (Mangin et al., 2026; Pickering et al., 2021), rather than passively redistributing resources carrying little cost. Distinguishing between these possibilities is critical because if facing pain is effortful, it may only be sustainable short term. Overtime, accumulated effort could lead to fatigue (Mangin & Pageaux, 2025), limiting long-term effectiveness, particularly in clinical populations.

When task demands remain within an individual’s capacity, performance may be preserved through compensatory resource mobilization. This position aligns with the motivational intensity theory (Brehm & Self, 1989), which conceptualizes effort as resource allocation in response to task demands and competing constraints. For submaximal tasks, increased difficulty, such as induced by pain, can be accommodated through greater effort investment (Silvestrini & Gendolla, 2019), thereby preserving performance. Within this framework, maintaining performance in the face of pain is inherently effortful.

A recent study from our group showed that individuals increased effort to maintain task performance under low and high pain during a visuomotor force-matching task and a cognitive interference task (Mangin et al., 2026). This preservation was not passive: effort increased with pain intensity, enabling task performance maintenance.

Building on these findings, we aim to identify neural alterations underlying this pain-related increase in effort. Using the same motor paradigm adapted for functional magnetic resonance imaging (fMRI), we investigate the neural mechanisms through which maintaining performance under pain becomes effortful.

Pain perception recruits well-established brain networks, including spino-thalamocortical pathways, somatosensory cortices, insula, anterior cingulate cortex, and supplementary motor area (Wager et al., 2013). By contrast, the neural substrates of effort perception remain poorly understood despite its central role in behavioral regulation and task performance (Inzlicht et al., 2018; Mangin & Pageaux, 2026; Preston & Wegner, 2009). In the cognitive domain, effort is typically linked to cognitive control and executive functions engagement (Shenhav et al., 2017), whereas in the physical domain its neural basis during task execution remains unresolved (Bergevin et al., 2023; Mangin & Pageaux, 2026). Converging evidence supports a central origin of effort perception, arising from centrifugal motor command processes rather than peripheral feedback (Bergevin et al., 2023; Mangin & Pageaux, 2026; Pageaux, 2016). These processes likely involve cortical regions upstream of primary motor cortex (M1), including the pre-supplementary motor area, supplementary motor area (SMA), anterior midcingulate cortex (aMCC), and insula (de Morree et al., 2012; Zénon et al., 2015).

The SMA and aMCC are strongly interconnected, contribute to the corticospinal tract, and connect with motor, parietal, insular, and prefrontal regions (Hoffstaedter et al., 2014; Nachev et al., 2008). The aMCC has been implicated in action cost evaluation (Shenhav et al., 2016), whereas the SMA supports cognitive control and action integration (Nachev et al., 2008). Moreover, increased SMA and aMCC activity has been associated with heat pain and motor processes (Misra & Coombes, 2015), suggesting that pain-related modulation of these regions may sustain effortful performance maintenance.

In this preregistered fMRI study, participants performed a visuomotor task under painful and non-painful conditions while reporting their perceived effort. We aimed to identify brain regions (i) engaged during the concurrent experience of pain and effort exertion, and (ii) associated with effort perception. Based on prior evidence linking SMA and aMCC to both pain and effort processing, we adopted a deductive approach focusing a priori on these regions of interest. We hypothesized (i) increased task-related activity in SMA and aMCC under pain, and (ii) an association between SMA–aMCC activity and perceived effort. We complemented this a priori approach with an inductive exploratory whole-brain analysis to identify other brain regions that may contribute to the effortful performance maintenance under pain.

## Materials and Methods

This study was preregistered on the Open Science Framework prior to data collection (registration link: https://osf.io/9w5mr). Behavioral and fMRI data are available at: https://zenodo.org/records/20445958.

### Participants

Forty-six healthy, right-handed young adults (24 females and 22 males; all right-handed; mean age 27.65 years; range 18–40 years) volunteered in this study. Exclusion criteria included any history of chronic pain, neurological or psychiatric disorders, major medical conditions (e.g., cancer or cardiovascular disease), musculoskeletal impairments affecting hand function, uncorrected visual deficits, and current use of psychotropic or analgesic medications. All participants provided written informed consent and received CAD 80 for participation. The study protocol was approved by the Research Ethics Committee of the Centre de Recherche de l’Institut Universitaire de Gériatrie de Montréal (CRIUGM).

Six participants were excluded prior to analysis: four due to excessive head motion during fMRI acquisition, and two due to an inability to differentiate between painful and non-painful thermal sensations despite individual calibration (i.e., reporting exclusively pain or exclusively warmth).

The final sample size used for data analysis included 40 participants (20 females, 20 males; all right-handed; mean age = 27.55 ± 4.56 years; range = 18–40 years). For two participants (IDs 12 and 14), data from the first run were excluded due to improper thermode placement, resulting in insufficient pain perception. For one participant (ID 5), the fourth run was excluded due to premature termination caused by discomfort in the scanner. All remaining data were included in the analyses.

### General procedure

The study involved a within-participant design. Participants visited the laboratory on two occasions. The first visit served as a calibration and familiarization session. During this session, thermal stimulation intensity was individually calibrated, maximal voluntary contraction (MVC) peak force was assessed, and participants were familiarized with the visuomotor force-matching task and the visual analogue scales (VAS) used to rate perceived effort and pain. These procedures are described in detail below. The second visit corresponded to the experimental session, **Fig 1**. Participants performed the visuomotor force-matching task inside the MRI scanner under both painful and non-painful thermal stimulation conditions.

**Fig 1:**
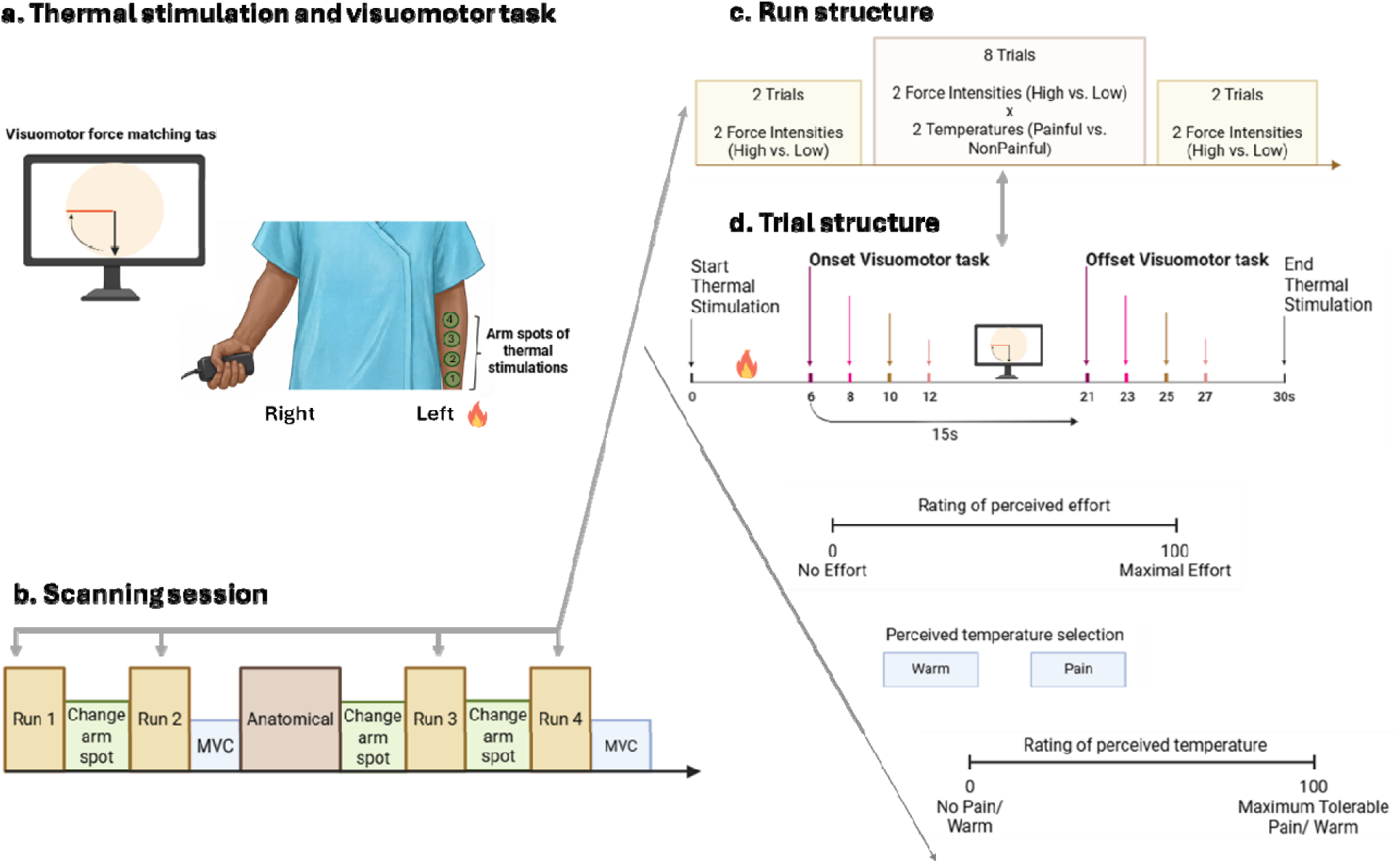
Overview of the experimental session. *Panel a* illustrates the visuomotor handgrip task performed with the right hand during thermal stimulation individually calibrated to elicit either high-pain (experimental condition) or non-painful warmth (control condition). Participants had to match the red line by squeezing the dynamometer at 5% (low force) or 30% (high force) of their maximal voluntary contractions (MVC) peak force. The illustration of a participant holding a dynamometer was created with the website https://conceptviz.app. *Panel b* illustrates the timeline of event in the scanner, including the acquisitions runs and anatomical scan, the change in arm spot for thermal stimulation to control for pain sensitization, and the completion of MVCs. *Panel c* shows the structure of each run. *Panel d* illustrates the structure of each trial. Each trial started with thermal stimulation (30 s duration) during which the visuomotor task was performed (15 s duration). The visuomotor task started randomly 6, 8, 10 or 12 after the onset of the thermal stimulation. Participants then rated the intensity of their perceived effort and the warmth/pain they experienced.

### Sensory calibration and thermal stimulations

Thermal stimuli were delivered using a 3 × 3cm MRI-compatible contact thermode applied to four skin sites on the volar surface of the left non-dominant forearm (TCS unit and T11 probe, QST.Lab, Strasbourg, France). Stimuli intensity ranged from 42°C (slightly warm) to a maximum of 50°C (max), with a plateau duration of 30 s. The maximum temperature of 50°C typically produces strong but tolerable pain and poses no risk of skin damage.

Painful thermal stimuli were individually calibrated in two steps. First, each participant’s pain tolerance temperature was estimated. Second, stimuli ranging from 42°C to the individual tolerance threshold were delivered to identify the temperature eliciting a target pain intensity of approximately 70/100 on a pain visual analog scale (VAS; 0_ = “no pain at all”, 100 = “maximum tolerable pain”). For all participants, the warm (non-painful) control temperature was set to 41.9°C. This temperature avoids the activation of nociceptors and reliably produces a non-painful warm sensation (Hallin et al., 1982; Van Hees & Gybels, 1981).

The individually calibrated painful temperature and the fixed warm temperature were subsequently used during the visuomotor force-matching task in the MRI scanner. Prior to the MRI scanning, participants were re-exposed to the target painful and warm temperature and asked to rate their perceptions on the VAS. To ensure consistency between sessions. If necessary, painful temperature was adjusted prior to the onset of the experimental task to maintain a target pain rating of approximately 70/100 on the VAS.

### MVC peak force calculation

The maximal voluntary contraction (MVC) peak force was collected via an fMRI-compatible handgrip dynamometer (TSD121B-MRI, BIOPAC, Goleta, USA). Maximal voluntary contractions were recorded during both the familiarization (Visit 1) and experimental (Visit 2) sessions. During Visit 1, MVCs were performed in a seated position with the right elbow at 90° and a right wrist brace to maintain the hand aligned with the forearm. During Visit 2, participants lied in the MRI machine. Consequently, MVCs were recorded in the supine position. The right wrist brace was again used to ensure alignment of the hand and forearm, with the upper limb fully extended. This position was maintained throughout the scanning session to standardize dynamometer positioning across trials and maximize force reproducibility.

The MVC peak force served as a reference value to set the target force during the visuomotor force-matching task. Before entering the MRI scanner, participants performed three 3 s MVCs, with 1-min rest in-between. The highest peak torque was used to set the low- and high- force of contraction intensities, respectively corresponding to 5% and 30% of MVC. To account for potential exercise induced decreased in force production capacity, additional MVCs were performed inside the scanner: one at the end of run 2, one at the end of the structural scan before run 3, and one at the end of run 4 (**Fig 1b**).

### Familiarization with the rating of perceived effort

During the first visit, participants were familiarized with the handgrip dynamometer and the VAS used to rate the intensity of perceived effort (0 = _“no effort”, 100 = _“maximal effort”). Participants performed a series of 10s force-matching contraction at the following intensities, in the following order 5%, 15%, 30%, 60%, 80%, 60%, 30%, 15%, 5% of their individual MVC peak torque. After each contraction participants rated the effort perceived to do the task, using the VAS. The perception of effort was defined as how hard, heavy, and strenuous it is to match and maintain the force feedback line (Marcora, 2010; Pageaux, 2016).

### Visuomotor force-matching task

The visuomotor force-matching task consisted of performing voluntary submaximal isometric handgrip contractions by squeezing a dynamometer to match a visually displayed force feedback line (**Fig 1a**).

Inside the MRI scanner, participants performed 15 s right isometric handgrip contractions at low- (5% MVC peak force measured outside the scanner) and high- (30% MVC peak force measured outside the scanner) force. Each contraction was followed by a pause to record perceived effort and perceived pain or warm, as well as to minimize the development of muscle fatigue (**Fig. 1d**). The perceptions of effort and pain or warm were assessed using the same VASs employed during the effort familiarization and sensory calibration procedures.

Force output was continuously recorded with the handgrip dynamometer, at a sample rate of 10 kHz, connected to a MP160 module (BIOPAC, Goleta, USA) and analyzed offline. Visual feedback of the force produced was provided on the screen using a circular force gauge. Participants were asked to align as precisely and steadily as possible the moving force signal with a force target line (baseline at 6 o’clock; target positioned at 9 o’clock) for both low- and high-force conditions. We decided to control for the anticipation of the force level condition by keeping the force target line at the same position on the screen. Therefore, more or less force was required to match the target force, depending on the high vs. low force condition The primary performance outcome was force steadiness, quantified with the coefficient of variation. Absolute and relative error were analyzed as secondary outcomes.

### Experimental Conditions and Procedure

The experimental session included four fMRI runs, with a total of 48 trials : 16 trials involved execution of the visuomotor task without any thermal stimulations (2 force levels × 8 trials); 16 trials involved execution of the visuomotor task during painful thermal stimulation (2 force levels × 8 trials); 16 trials involved execution of the visuomotor task during non-painful warm stimulation (2 force levels × 8 trials).

Each run included 12 trials presented in the following sequence: 2 trials of task execution at low- and high-force without thermal stimulation, 8 trials of task execution at low- and high-force with thermal stimulation and 2 trials of task execution at low- and high-force without thermal stimulation (**Fig 1c**).

For trials involving thermal stimulation, the onset of the trial coincided with the onset of the thermal stimulus, which lasted 30.5 s. The visuomotor contraction began 6, 8, 10, or 12 s after thermal onset (**Fig 1d**). This temporal offset avoided the simultaneous onset and offset of the thermal stimulus and motor task. For trials without thermal stimulation, contractions began 6, 8, 10, or 12 s after trial onset.

Trials without thermal stimulation served as control conditions for the visuomotor task, allowing identification of visuomotor task-related brain activations independent of thermal stimuli. The force level order of these trials was alternated between runs (e.g., run 1 started with contraction at low-force, run 2 with contraction at high-force). Following these contractions, participants only rated their perceived effort only as no thermal stimulation was applied.

The 32 trials performed during painful and warm stimulation were pseudo-randomized within and between runs, and counterbalanced across participants.

During the recovery period following each contraction under thermal stimulation, participants first rated the intensity of their perceived effort with a VAS. Then, they indicated whether the thermal stimulation was experienced as painful or warm, by choosing between two buttons presented on the screen and labelled “pain” and “warm”. Then, participants rated the intensity of pain or warmth on a VAS.

At the end of each run, the experimenter moved the thermal stimulator to a different arm spot (4 different arm spots were used) to prevent a possible sensitization of the stimulated area.

### Questionnaires

The Pain Catastrophizing Scale (PCS) (Sullivan et al., 1995) was administered once during the familiarization session to assess participants’ tendency to magnify or ruminate about pain.

At the beginning and end of the experimental session, participants completed the VAS assessing subjective fatigue, boredom, and motivation (confounds of effort perception (Mangin & Pageaux, 2025)). Results of these VAS are presented in Supplementary Information S6.

Fatigue was measured on a scale ranging from 0 (“no fatigue at all”) to 100 (“extremely fatigued”) and assessed subjective feelings of fatigue. Boredom was rated from 0 (“not bored at all”) to 100 (“extremely bored”) and assessed perceived boredom during the session. Motivation was assessed before and after the scan. Before scanning, the scale measured motivation to complete the session. After scanning, the scale measured the willingness to repeat the session. Both ranged from 0 (“no motivation at all”) to 100 (“extremely motivated”).

### Statistical analysis

#### Behavioral data analysis

Behavioral data were analysed using Jamovi (version 2.3.21) and R. All analyses were converted into R scripts that have been shared on Github (https://github.com/me-pic/MRI_pain_effort). Fatigue, motivation and boredom scores were analysed using paired-sample t-test comparing pre- and post-MRI measurements.

Ratings of perceived effort were analysed using linear mixed-effects model. Stimulation temperature (warm vs painful) and contraction force (low vs high) were entered as categorical fixed effects (factors), and participant was included as a random effect. A similar model was used to analyse ratings of perceived temperature, with warm and pain ratings considered from 0 to 200. A score below 100 indicates a warm, non-painful sensation, and a score above 100 indicates a painful sensation. For painful ratings, the score was added to a constant *k* = 100 to obtain a score > 100.

#### Force analysis

The force signal was extracted using LabChart 8 software (AD Instrument, USA). For each contraction, raw signal was analysed. The mean force and the standard deviation (SD) were calculated over the last 5 seconds of the contraction plateau. To do so, we calculated the last 6 seconds of contraction starting from the offset of each contraction, identified as the time point where force starts decreasing to reach the baseline value. Then we subtracted 1 s, to ensure that we were at a plateau, and then used the 5 remaining seconds. Subsequently, we calculated the coefficient of variation to estimate whether the force variability remained steady across conditions. The coefficient of variation was our primary performance outcome. We also calculated as secondary performance outcome the absolute and relative error.

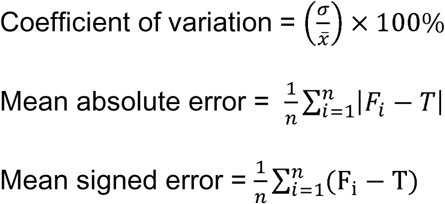

where σ is the standard deviation, x is the mean, *F*_i_ is the force exerted by the participants, and *T* is the target force. Finally, a statistical analysis was performed using a linear mixed-effects model in R (version 4.3.3), with stimulation temperature (non-painful vs. painful) and force level (low vs. high) as fixed effect, and participant as a random effect.

#### fMRI acquisition

Imaging data was acquired on a 3T Siemens Prisma Fit scanner equipped with a 32-channel head coil. Participants’ heads were stabilized in a comfortable position using a vacuum bag. Participants were instructed to as still as possible throughout the scanning session and wore earplugs to attenuate scanner noise. Each participant underwent a T1-weighted anatomical scan using a ME-MPRAGE sequence, between the second and third functional runs (TR: 2200 ms; TE: 1.87 ms; flip angle: 8°; voxel size: 1 × 1 × 1 mm; acquisition matrix: 256 × 256; number of slices: 176). The functional scans were collected using a blood oxygen level-dependent (BOLD) protocol with a T2_∗_-weighted gradient echo-planar imaging sequence (Multiband acceleration factor: 3; TR: 0.832 s; TE: 20 ms; flip angle: 70°; voxel size: 3 × 3 × 3 mm; acquisition matrix: 64 × 64; number of slices: 51).

#### fMRI preprocessing

Data were preprocessed using fMRIPrep (version 23.2.1)(Esteban et al., 2019). Preprocessing steps included a head-motion correction, a co-registration between the functional and anatomical images, and the resampling of the preprocessed BOLD run in a standard space (MNI152NLin2009cAsym). The framewise displacement (FD) was computed and used as an exclusion criterion. Frames that exceeded a threshold of 0.5 mm FD or 2 standardized DVARS were annotated as motion outliers. Four participants excluded, see subsection “Participants” of this Methods section. For more details about the preprocessing procedure, see Supplementary Information S7. Furthermore, the preprocessed BOLD images were smoothed prior to the GLM with 6mm FWHM Gaussian Kernel.

#### fMRI analyses

All fMRI analyses were conducted using custom Python scripts (Python version 3.9.23) using the Nilearn package (version 0.10.3). All scripts are available on Github (https://github.com/me-pic/MRI_pain_effort). An overview of fMRI analysis procedures is presented in **Fig 2**. Brain activation analyses included a hypothesis-driven approach focusing on the SMA and aMCC (FDR-corrected, *q* < 0.05), complemented by an exploratory whole-brain analysis (FDR-corrected, *q* < 0.01).

**Fig 2:**
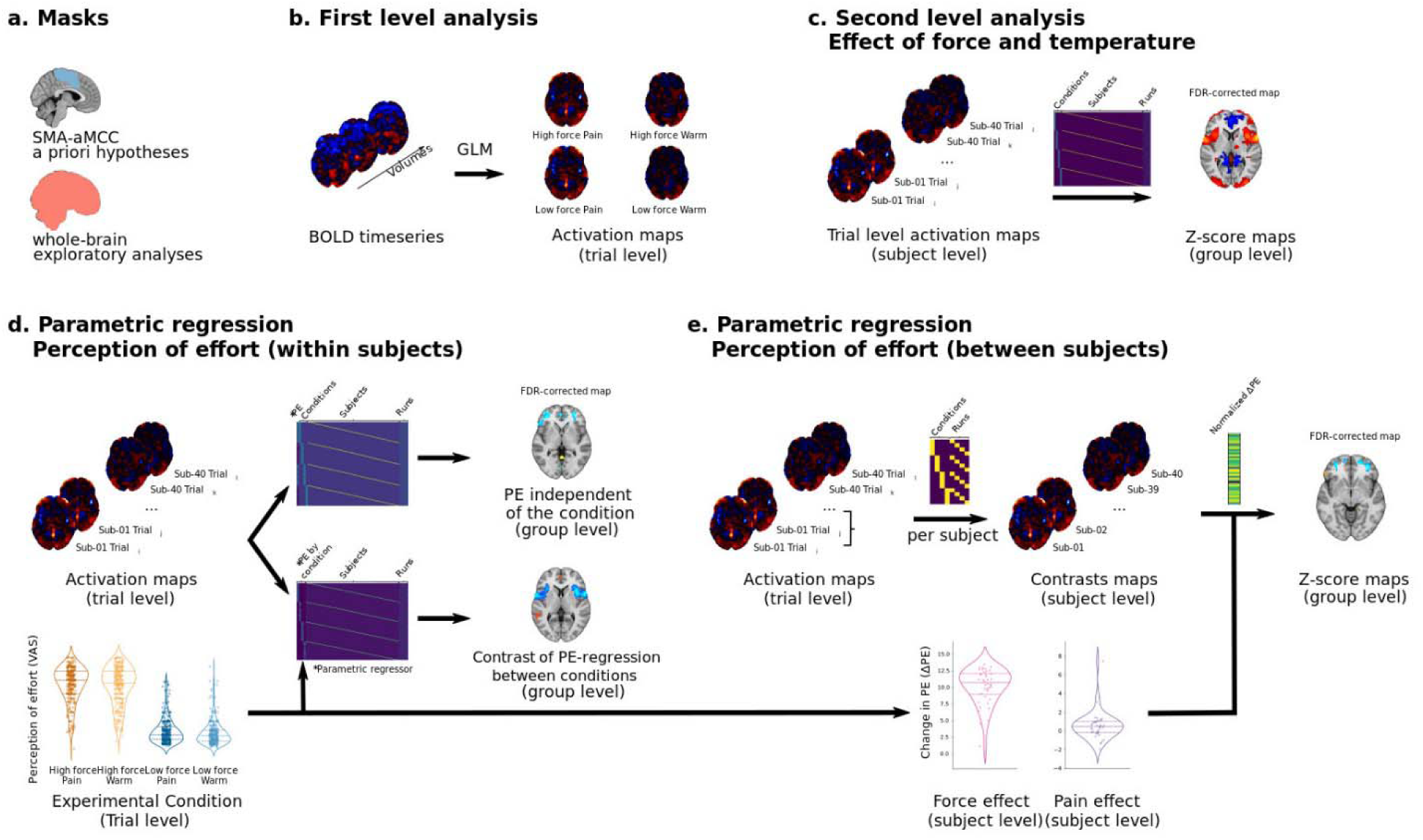
Overview of the fMRI pipeline analysis. *Panel a*: Masks used for the fMRI analyses. Primary analyses were conducted at the voxel level within a priori regions of interest SMA-aMCC using false discovery rate (FDR) correction at q<.05. Exploratory analyses were further performed on the whole-brain using FDR correction at q<.01. *Panel b*: For each participant, trial-by-trial activation maps were generated from the preprocessed BOLD timeseries. *Panel c:* These trial-by-trial maps were entered into group level analyses to identify brain regions associated with execution of the visuomotor task under high-force and pain conditions. *Panel d:* To obtain brain regions related to perception of effort (PE), trial-by-trial maps were incorporated into group level parametric regression analyses. We first identified brain regions correlating with PE ratings independently of the experimental condition (top matrix), and then we identified brain regions correlating with PE ratings differentially as a function of force level or thermal stimulation (bottom matrix). *Panel e:* To evaluate brain areas related with interindividual differences in PE, individual-level contrast maps were computed for the contrasts of interest: Contraction_High (Painful + NonPainful) > Contraction_Low (Painful + NonPainful) and Contraction_Painful (High + Low Force) > Contraction_NonPainful (High + Low Force). These maps were then entered into a parametric regression as dependent variable, and the normalized delta PE ratings were used as independent variable.

The general linear model (GLM) was used to quantify single-trial response magnitudes at the run level for each participant. The GLM design matrix was constructed using separate regressors for each trial. First, boxcar regressors convolved with the canonical hemodynamic response function were constructed to model thermal stimulation and task execution for each trial and run. Then, confounds related to cerebral blood flow (2 covariates), white matter (2 covariates), motion artifacts (24 covariates), and low-frequency noise (2 covariates) were entered in the design matrix, for a total of 30 nuisance regressors. Multiple participant-level GLMs were performed to answer different questions. Details are described in the sections below.

#### Brain masks

Since we had hypotheses on a priori regions, we computed a SMA-aMCC mask using the Schaefer atlas with 100 region of interests (ROIs) (https://nilearn.github.io/dev/modules/description/schaefer_2018.html). The parcellations aligned with 2mm FSL MNI template was used. The ROIs with the following labels were included in the mask: “7Networks_RH_SalVentAttn_Med_2”, “7Networks_RH_SomMot_8”, “7Networks_LH_Cont_Cing_1”, “7Networks_LH_SalVentAttn_Med_3”, “7Networks_LH_SalVentAttn_Med_1”, “7Networks_LH_SomMot6”.

The analyses were also performed at the whole-brain level (group mask obtained from the T1W individual images). All analyses were first performed using the SMA-aMCC mask (a priori hypothesis), then using the whole-brain mask. Results are presented as such.

#### Manipulation check – Pain-related brain activation at rest

To verify the sensitivity of our protocol in detecting pain-related brain activity, we modelled the thermal trials (Painful and Non-painful stimulation) from the beginning of the thermal stimulation to the beginning of the motor contraction in a first level GLM. By doing so, we isolated brain responses associated with thermal processing, excluding motor-related activity.

The obtained individual trial-by-trial activation maps were concatenated in a group level GLM in which participants and runs regressors were included as covariates. The Painful > NonPainful contrast was computed at the group-level.

#### Manipulation check – Visuomotor task execution-related brain activation without thermal stimulation

To ensure that we were able to detect motor-related brain activation independently of thermal stimulations, we performed a participant-level GLM on the contraction trials without thermal stimulation.

For each run, we modelled trials executed at low-force (ContractionSolo_Low) and high-force (ContractionSolo_High). We then concatenated the trial-by-trial activation maps across runs and participants in a group-level GLM in which two regressors were used to model the two force levels and confounding regressors were added for runs and participants. Paired t-tests were performed on each regressor of interest to make sure that motor-related areas were observed at both forces. The contrast ContractionSolo_High > ContractionSolo_Low was also computed.

#### Identifying brain areas related to the execution of the visuomotor task in the presence of pain

To address our objectives, we performed a participant-level GLM to model the trials in which the thermal stimulation and the motor contractions were performed concurrently. The following trial-by-trial activation maps were obtained for each run and each participant: Contraction_High_Painful (trials executed at high-force during painful stimulation), Contraction_Low_Painful (trials executed at low-force during painful stimulation), Contraction_High_NonPainful (trials executed at high-force during non-painful stimulation), Contraction_Low_NonPainful (trials executed at low-force during non-painful stimulation).

Using the trial-by-trial activations, a group-level analysis was performed including the experimental regressors (Contraction_High_Painful, Contraction_Low_Painful, Contraction_High_NonPainful and Contraction_Low_NonPainful) and the confounding regressors (one regressor per participant and one regressor per run). To test the effect of the motor contractions in the presence of thermal stimulation, a voxelwise paired t-test was computed for the contrast Contraction_High (Painful + NonPainful) >

Contraction_Low (Painful + NonPainful). To obtain the brain areas related to the effect of pain in presence of the motor contractions, a paired t-test was performed on the contrast Contraction_Painful (High + Low) > Contraction_NonPainful (High + Low).

#### Identifying brain areas related to the perception of effort

*Within-participants – Across conditions*.

To identify within-participant brain areas related to spontaneous fluctuations in perception of effort, we first performed a group-level parametric regression concatenating the trial-by-trial activation maps across all conditions (i.e., Contraction_High_Painful, Contraction_Low_Painful, Contraction_High_NonPainful and Contraction_Low_NonPainful) across subjects. The perception of effort was modelled as one continuous regressor, while subject- and run-specific effects were included as confounding variables. A paired t-test was performed on the parametric regressor slope to identify the voxels for which the activation correlated significantly with the perception of effort independently of any condition.

*Within-participants – By conditions*.

To assess whether the relationship between the perception of effort and brain activity differed across experimental conditions within participants, we performed another group-level parametric regression using the same trial-by-trial maps but now including the perception of effort as separate regressors for each condition. From this model, two voxel-wise paired t-tests were computed: i) Contraction_High (Painful + NonPainful) > Contraction_Low (Painful + NonPainful) and ii) Contraction_Painful (High + Low) > Contraction_NonPainful (High + Low).

#### Between-participants

To investigate the brain areas that covaries with the perception of effort across participants, a between–participant parametric regression analysis was conducted. For each participant, contrasts of interest (i.e., Contraction_High vs Contraction_Low regardless of the thermal intensity, and Contraction_Painful vs Contraction_NonPainful regardless of the motor force intensity) were first computed on both the fMRI and behavioural data. The resulting participant-level activation maps were then integrated in separate group-level regression models (one model per contrast) in which normalized PE ratings served as the only regressor. The PE ratings were normalized using a min–max normalization procedure according to the following formula: (delta PE - delta PE_min) / (delta PE_max - delta PE_min).

Delta PE was computed as the difference between perceived effort during high-force and low-force (PE ratings_Contraction_High – PE ratings_Contraction_Low). Delta PE_min and delta PE_max correspond to the minimum and maximum delta PE values observed across participants. One sample t-test on the regressor slope was performed at the voxel level to identify regions where the brain activity correlated with interindividual differences in perceived effort.

#### Interpretation of Parametric Regressions

Because both force level and pain modulated perceived effort, parametric regression analyses were conducted to identify brain regions whose activity covaried with trial-by-trial perceived effort ratings. The interpretation of these regressions depended on the direction of the task-related BOLD response (activation vs. deactivation relative to baseline) and the sign of the parametric regression coefficient (β). Four scenarios were considered:

1. Positive task-related activation (BOLD > 0) and positive regression coefficient (β > 0). This pattern indicates that greater perceived effort was associated with increased task-related activation.
2. Negative task-related activation (BOLD < 0) and negative regression coefficient (β < 0). This pattern indicates that greater perceived effort was associated with stronger task-related deactivation.
3. Positive task-related activation (BOLD > 0) and negative regression coefficient (β < 0). This pattern indicates that greater perceived effort was associated with reduced task-related activation.
4. Negative task-related activation (BOLD < 0) and positive regression coefficient (β > 0). This pattern indicates that greater perceived effort was associated with reduced task-related deactivation.

This framework allowed us to distinguish whether effort ratings were associated with amplification, suppression, or attenuation of task-related neural responses.

## Results

Participants executed, in the presence or absence of thermal stimulation, submaximal isometric handgrip contractions to match as precisely as possible a visual target at two force levels (5% and 30% of maximal voluntary peak force assessed outside the scanner), corresponding to 10.1 ± 2.2% and 51.9 ± 9.3% of maximal force exerted in the scanner, respectively.

### Behavioral results

Results in the text are presented as mean ± standard deviation.

#### Pain induction

Thermal stimulation successfully induced pain. Ratings increased with temperature from 27.3 ± 15.4 (non-painful warmth; <100) to 162.7 ± 17.4 (painful; >100), revealing a robust main effect of temperature, F(1, 39.2) = 1306.21, *p* < .001. There was a temperature × force interaction [F(1, 8.07) = 1176.9, *p* = .005]. In the warm temperature condition, participants reported lower warmth perception at low force (21.9 ± 12.4) compared to high force (32.6 ± 21.5) condition (t(1177) = −4.798, *p* < .001). In the pain temperature condition, the force level did not influence the pain rating (low force 161.9 ± 18.6; high force 163.7 ± 17.4; t(1176.9) = −0.798, *p* > .999).

#### Performance

Participants were instructed to match the force feedback line as precisely and steadily as possible. Force steadiness, quantified as the coefficient of variation, served as the primary performance outcome (**Fig 3a**). Force steadiness was not affected by temperature [F(1, 78) = 0.24 *p* = .624], indicating comparable performance under painful and non-painful conditions. Steadiness was higher during low- compared to high-force contractions [F(1, 39) = 115.27, *p* < .001], with no temperature × force interaction [F(1, 78) = 0.16, *p* = .691]. Mean absolute and signed error were analyzed as secondary outcomes (Supplementary Information S1) and yielded a similar pattern, confirming preserved performance under pain.

**Fig 3:**
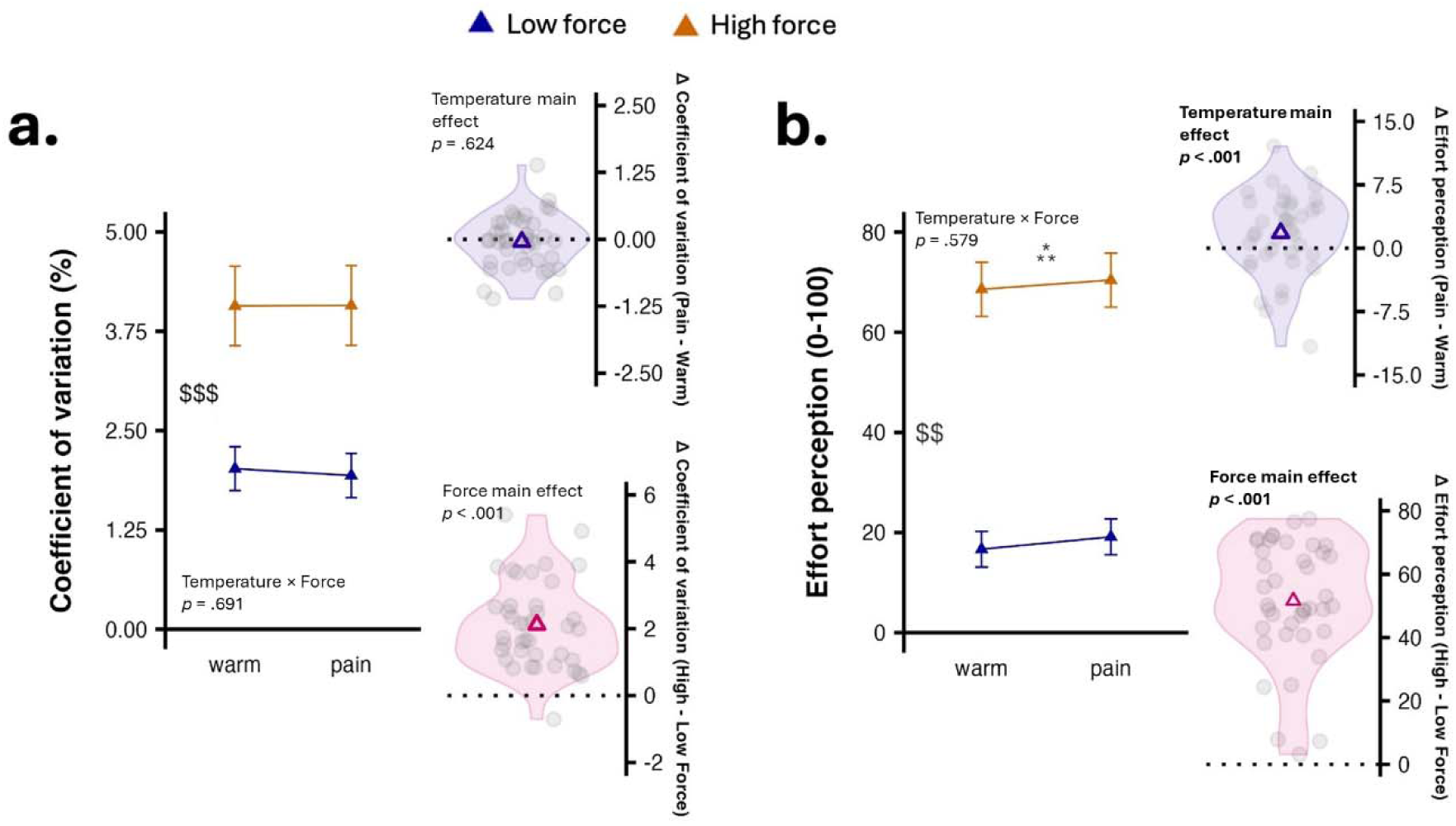
Performance and perception of effort during the visuomotor task under pain and warm condition. *Panel a*: performance was measured as force steadiness, inferred from the coefficient of variation. *Panel b*: Participants rated their perceived effort with a visual analog scale. The visuomotor task was performed for 15 s at low and high force levels, corresponding to 5% and 30% of individual maximal voluntary force measured outside the scanner. * and ^$^ represents a main effect of thermal stimulation and force level, respectively. Two and three symbols for *p* < .01 and *p* < .001, respectively. Data is presented as mean ± 95%CI.

#### Perception of effort

Perceived effort ratings are shown in **Fig 3b**. Performing the visuomotor task from low to high force increased perceived effort [F(1, 39) = 293.87, p < .001]. Effort ratings were also higher under painful compared to non-painful stimulation [F(1, 1176.3) = 12.16, p < .001], with no temperature × force interaction [F(1, 1176.3) = .307, p = .579].

Together, these findings indicate that performance was maintained under pain at both force levels, while perceived effort increased.

#### Brain activation

Brain activation analyses included a hypothesis-driven approach focusing on the SMA and aMCC (FDR-corrected, *q* < 0.05), complemented by an exploratory whole-brain analysis (FDR-corrected, *q* < 0.01).

To separate pain-related activity from motor processes, we modelled thermal stimulation from its onset to the start of the visuomotor task. The contrast between painful and non-painful stimulation revealed activation patterns consistent with established pain-related networks (Wager et al., 2013). To isolate force-related activity independently of thermal effects, we modelled task execution at both force levels during trials without thermal stimulation. The high- versus low-force contrast identified activation patterns consistent with established motor and sensorimotor networks (Dai et al., 2001; Vaillancourt et al., 2003). Suppression of default mode network activity during task engagement (Raichle et al., 2001) with increasing task demands (Čeko et al., 2015; Weber et al., 2022), as well as reduced basal ganglia activity at higher force levels (Spraker et al., 2007; Vaillancourt et al., 2003), aligned with prior findings. Details are provided in Supplementary Information 2 and 3.

#### Execution of the visuomotor task in the presence of pain

To identify regions engaged during concurrent pain and effort exertion, we modelled task execution under painful and non-painful thermal stimulation, independently of the force level, and contrasted the two conditions. See Supplementary Information 4 for the separate brain activation maps related to the task execution under painful and non-painful thermal stimulation. For the contrast, a priori ROI analysis (**Fig 4a**) revealed reduced activation under pain in bilateral SMA, as well as stronger deactivation in midcingulate regions, including left MCC and right aMCC. Exploratory whole-brain analysis (**Fig 4b**) showed increased activation under pain in ventral orbitofrontal cortex (OFC), subgenual and dorsal ACC, posterior cingulate cortex (PCC), precuneus, lateral prefrontal cortex, superior frontal gyrus, paracentral lobule, angular gyrus, and middle temporal cortex. Decreased activation was observed in bilateral anterior insula, pre- and postcentral gyri, SMA/MCC, parietal and frontal opercula, and cerebellum.

**Fig 4:**
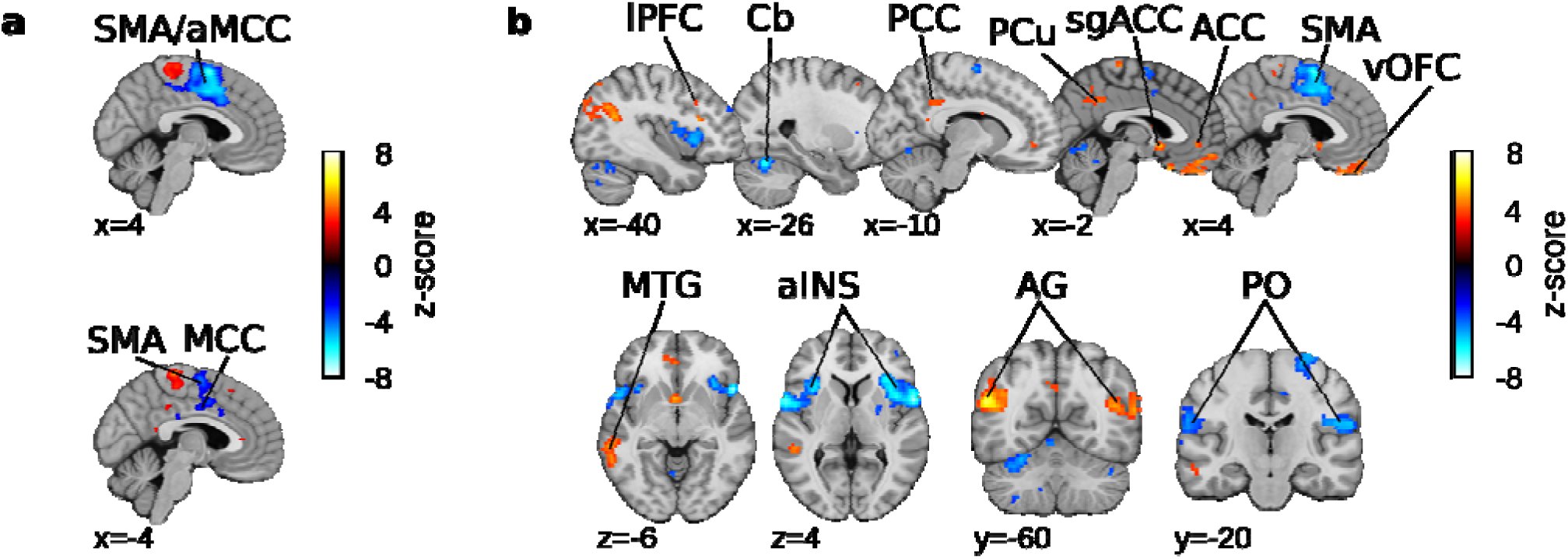
Brain areas associated with visuomotor task execution during painful stimulation. Activation maps presenting the contrast between the visuomotor task executed during painful stimulation and warm non painful stimulation (task execution during pain > task execution during warm contrast), independently of the force level. *Panel a*: Z-score map computed on the SMA-aMCC mask (q<0.05; FDR corrected; a priori hypothesis focused on regions of interest). *Panels b*: Z-score map computed at the whole-brain level (q<0.01; FDR corrected; exploratory whole brain analysis). AG, angular gyrus; aINS, anterior insula; aMCC, anterior midcingulate cortex; Cb, cerebellum; lPFC, lateral prefrontal cortex; MCC, midcingulate cortex; MTG, middle temporal gyrus; OFC, orbitofrontal cortex; PCC, posterior cingulate cortex; PCu, precuneus; PO/SII, parietal operculum/secondary somatosensory cortex; SMA, supplementary motor area; (sg)ACC, (subgenual) anterior cingulate cortex.

#### Brain areas related to the perception of effort

Because both force level and pain modulated perceived effort, we performed parametric regression analyses to identify brain regions whose activity covaried with effort ratings. Interpretation of these effects considered the direction of task-related activation (activation vs. deactivation) and the sign of the regression coefficients, allowing us to distinguish between increased activation, increased deactivation, or attenuation of task-related responses as perceived effort rose.

#### Brain areas related to the perception of effort, independently of pain or force level – Within-participants analysis

We were interested in identifying voxels whose activation correlated with perceived effort ratings, independently of the experimental manipulations (increasing force or pain). We conducted a within-participants parametric regression analysis where we regressed trial-by-trial activation maps with trial-by-trial perceived effort ratings, while controlling for the differences between conditions (covariance).

A priori approach (**Fig 5a**) revealed some voxels displaying a positive correlation in left SMA and a negative correlation in bilateral MCC. Exploratory whole brain analysis (**Fig 5b**) revealed additional positive correlations with bilateral post-central gyrus, right precuneus, right cuneus, right middle temporal gyrus and cerebellum; and negative correlation with bilateral medial prefrontal cortex (medial PFC), ventromedial prefrontal cortex (vmPFC), lateral prefrontal cortex (lateral PFC), lateral orbitofrontal cortex (lateral OFC), left PCC, right ventral striatum, left middle temporal gyrus.

**Fig 5:**
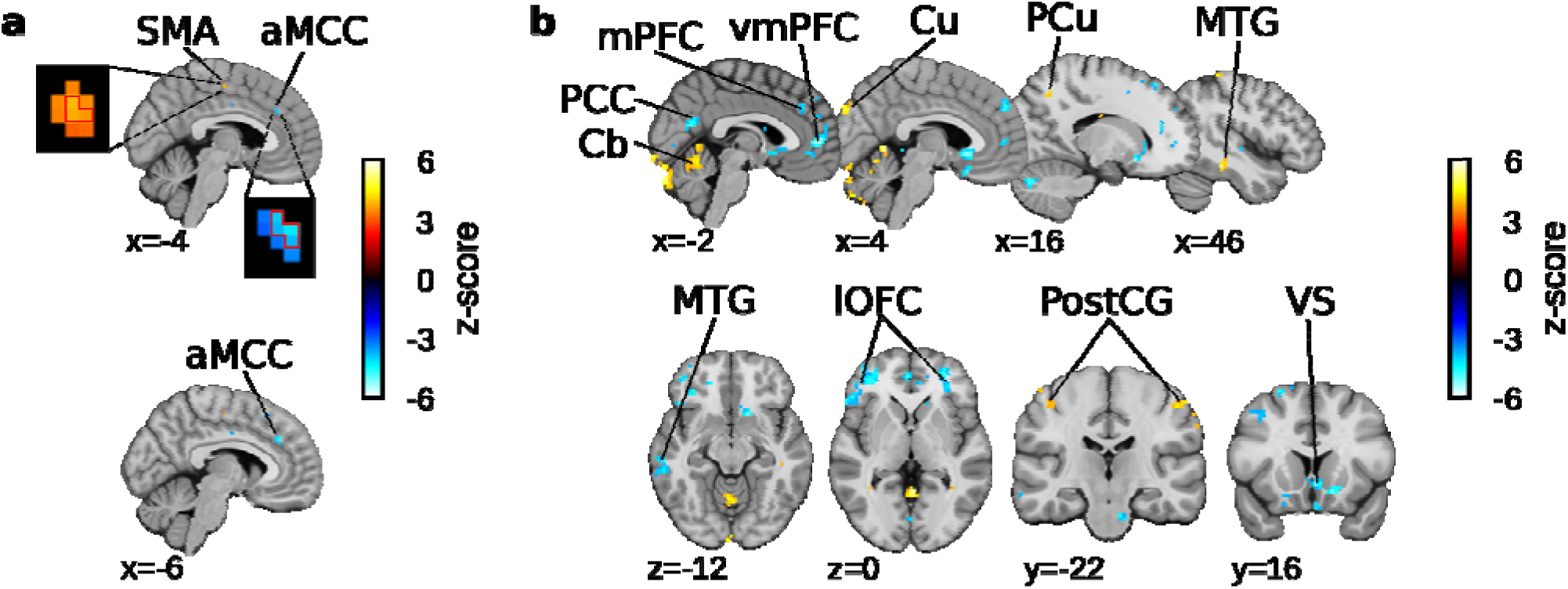
Brain areas associated with effort perception, regardless of the experimental condition, within-participants parametric regressions*. Panel a*: Z-score map computed on the SMA-aMCC mask (q<0.05; FDR corrected; a priori hypothesis focused on regions of interest). Insets show the activated voxels at *p*-uncorrected < 0.001, with red-outlined voxels significant at FDR-corrected q<0.05. *Panels b*: Z-score map computed at the whole-brain level (q<0.01; FDR corrected; exploratory whole brain analysis). aMCC, anterior midcingulate cortex; Cb, cerebellum; Cu, cuneus; lPFC, lateral prefrontal cortex; mPFC, medial prefrontal cortex; MTG, middle temporal gyrus; PCC, posterior cingulate cortex; PCu, precuneus; PostCG, post central gyrus; SMA, supplementary motor area; VS, ventral striatum.

#### Brain areas related to the perception of effort, increased force level and pain – Within-participants analysis

To assess whether the relationship between the perception of effort and brain activity differed within participants across experimental conditions, we performed voxel-wise regression analyses. Activation maps at a trial level were used as dependent variables and perceived effort ratings as predictors. Then, the regression maps obtained in the different condition were compared using the following contrasts: high versus low force (collapsed across thermal stimulation) and painful versus warm stimulation (collapsed across force levels). This approach allowed us to identify brain regions in which brain activity scaled with perceived effort differently across force and thermal stimulation conditions.

#### Increasing force and perceived effort

A priori approach (**Fig 6a**) revealed positive correlation with bilateral SMA and negative correlation with bilateral aMCC. Exploratory whole brain analysis (**Fig 6b**) revealed positive correlations with left and right pre-central gyrus, bilateral SMA, left frontal operculum and cerebellum; and negative correlation with bilateral ACC and vmPFC, aMCC, bilateral superior temporal gyrus, bilateral posterior insula and caudate nucleus.

**Fig 6:**
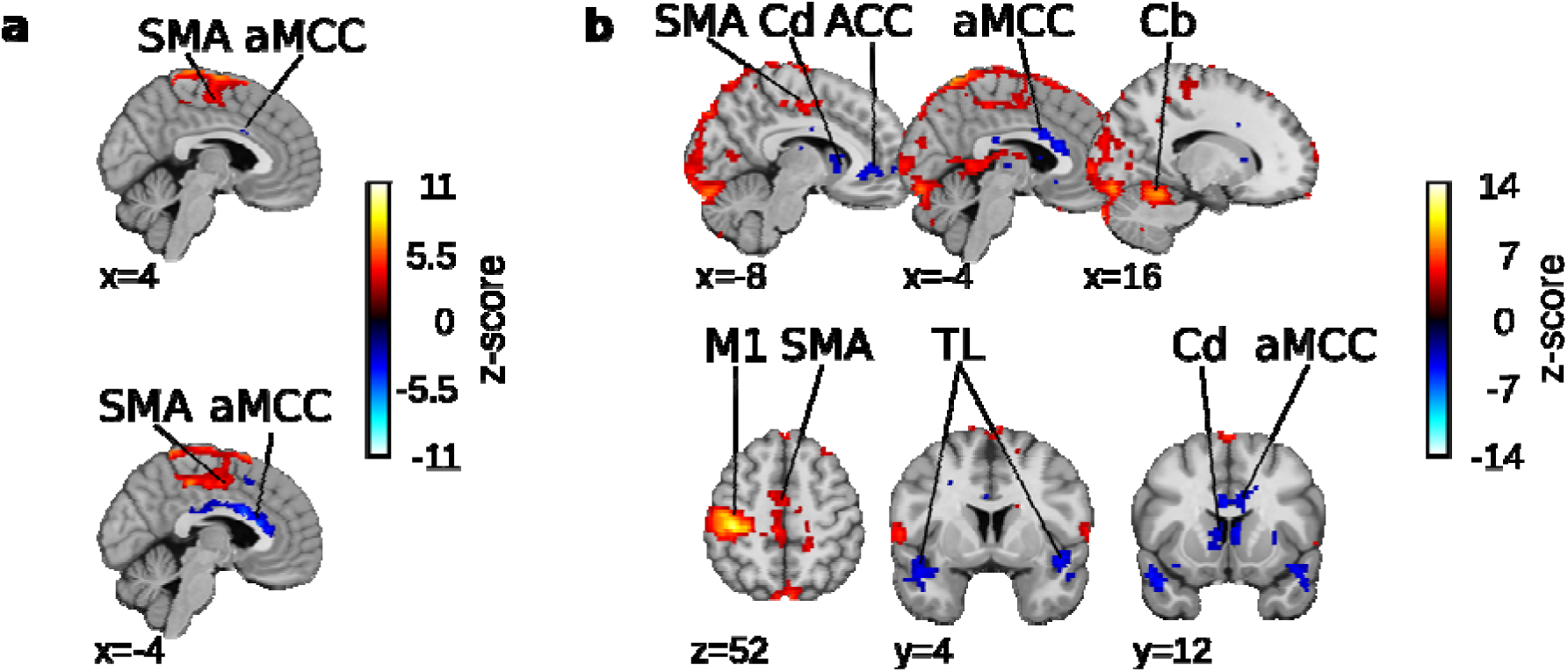
**Brain areas associated with effort perception and increased force level of the visuomotor task, within-participants parametric regressions**. *Panel a*: Z-score map computed on the SMA-aMCC mask (q<0.05; FDR corrected; a priori hypothesis focused on regions of interest). *Panel b*: Z-score map computed at the whole-brain level (q<0.01; FDR corrected; exploratory whole brain analysis). ACC, anterior cingulate cortex; aMCC, anterior midcingulate cortex; Cb, cerebellum; Cd, caudate nucleus; M1, primary motor cortex; SMA, supplementary motor area; TL, temporal lobe.

#### Pain effect and perceived effort

A priori approach (**Fig 7a**) revealed negative correlations with right SMA and bilateral aMCC. Exploratory whole brain analysis (**Fig 7b**) revealed positive correlations with bilateral ventral OFC, subgenual ACC, precuneus, angular gyrus, middle temporal gyrus and superior frontal gyrus, left dorsolateral prefrontal cortex (DLPFC) and PCC; and negative correlations with bilateral anterior insula and frontal operculum, right SMA and aMCC and right DLPFC.

**Fig 7:**
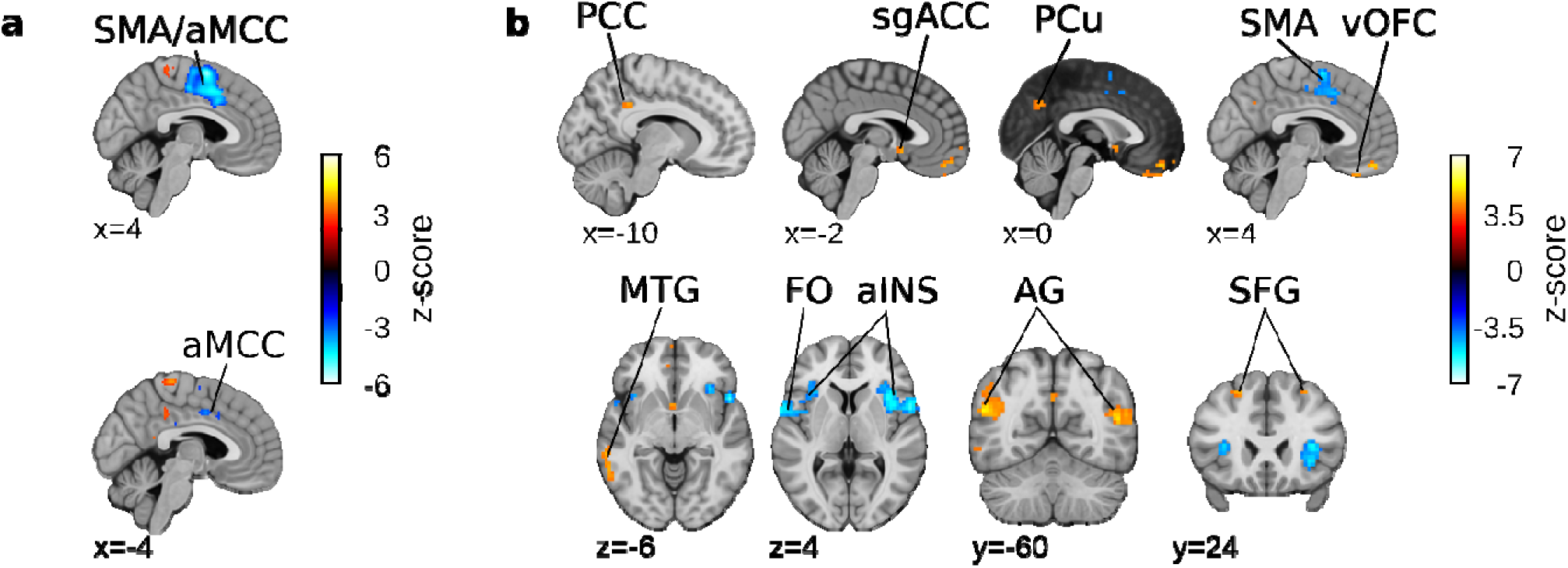
**Brain areas associated with effort perception and pain during the visuomotor task, within-participants parametric regressions**. *Panel a*: Z-score map computed on the SMA-aMCC mask (q<0.05; FDR corrected; a priori hypothesis focused on regions of interest). Panel b: Z-score map computed at the whole-brain level (q<0.01; FDR corrected; exploratory whole brain analysis). AG, angular gyrus; aINS, anterior insula; aMCC, anterior midcingulate cortex; FO, frontal operculum; MTG, middle temporal gyrus; PCC, posterior cingulate cortex; PCu, precuneus; SFG, superior frontal gyrus; sgACC, subgenual anterior cingulate cortex; SMA, supplementary motor area; vOFC, ventral orbitofrontal cortex.

#### Brain areas related to the perception of effort – Between-participants analysis

We conducted between–participant parametric regression analyses to identify brain regions which activity correlated with interindividual differences in perceived effort. Using the contrast high versus low force (collapsed across thermal stimulation), our a priori approach (**Fig 8a**) revealed positive correlation with left SMA and negative correlation with left MCC and right aMCC. Exploratory whole brain analysis (**Fig 8b**) revealed positive correlation with left and right pre-central gyrus, left SMA, and cerebellum; and negative correlation with ACC and medial PFC, MCC, lateral OFC, insula, thalamus and basal ganglia. Using the contrast painful versus warm stimulation (collapsed across force levels), our a priori approach did not reveal significant effects in the SMA and aMCC after FDR correction. Exploratory whole brain analysis revealed positive correlations with right paracentral lobule, bilateral ventral OFC, left vmPFC, left IFG (pars triangularis), right temporal pole; and negative correlation with left and right lateral OFC, right superior parietal lobule, left middle temporal cortex, right putamen and cerebellum (see Supplementary Information S5).

**Fig 8:**
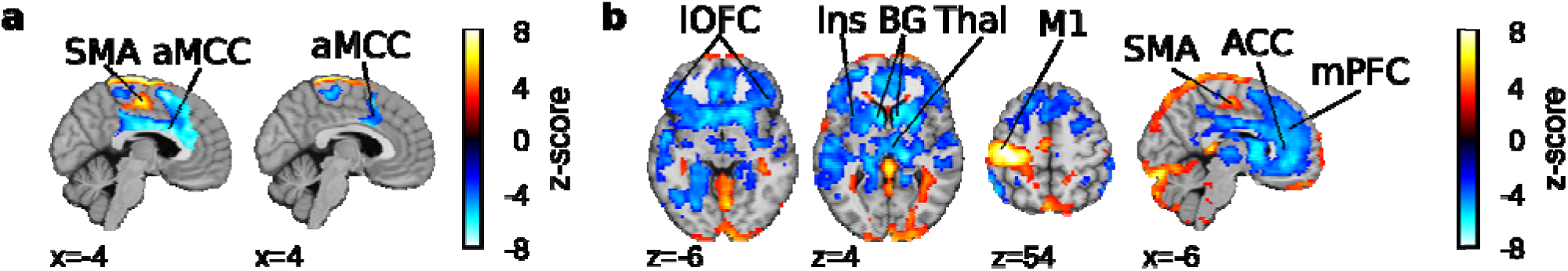
Brain area associated with effort perception and increased force level of the visuomotor task. Between participants parametric regressions were performed. *Panel a*: Z-score map computed on the SMA-aMCC mask (q<0.05; FDR corrected; a priori hypothesis focused on regions of interest). *Panel b*: Z-score map computed at the whole-brain level (q<0.01; FDR corrected; exploratory whole brain analysis). aMCC, anterior midcingulate cortex; BG, basal ganglia; Ins, insula; lOFC, lateral orbitofrontal cortex; M1, primary motor cortex/pre-central gyrus; mPFC, median prefrontal cortex; SMA, supplementary motor area; Thal, thalamus.

## Discussion

In this study, we show that maintaining visuomotor performance under thermal pain is possible but accompanied by increased perceived effort. The dissociation between preserved task performance and elevated perceived effort indicates that successful task execution under pain is not cost-free. Rather than reflecting passive attentional redistribution, task performance maintenance appears to rely on active resources allocation. Consistent with motivational intensity theory (Richter et al., 2016), effort may reflect additional resources mobilized to counteract pain-related interference while keeping task performance stable. Although this behavioral adaptation may remain effective short term, it may become detrimental overtime as fatigue results from accumulating effort (Van Damme et al., 2018).

Pain without task execution (at rest) induced activation patterns consistent with established pain-related networks (Wager et al., 2013), including increased SMA and aMCC activity. Executing the visuomotor force-matching task under painful and non-painful stimulation increased SMA activity and deactivated aMCC. Contrary to our hypothesis that pain would enhance activity in these regions, the pain vs. warm contrast during task execution and the activation maps of painful and non-painful task execution, revealed reduced SMA activity and stronger aMCC deactivation. This finding differs from Misra and Coombes (2015), who reported increased aMCC activity during force-matching under pain. This discrepancy likely reflects differences in trial structure. In their study, painful stimulus onset was simultaneous with the visuomotor task, and trials were not randomized between conditions (changing every six trials), allowing participants to anticipate the upcoming painful or non-painful trials. In our study, painful stimulation always preceded task onset by several seconds, and trials were randomized between condition, controlling for pain anticipation. Given that pain expectation modulates brain activity in pain processing regions (Atlas & Wager, 2012), the absence of temporal separation between thermal stimulus onset and task execution in their design likely contributed to anticipatory effects, which may explain the discrepancy between findings. Future studies should directly test how pain anticipation prior to task onset influences brain activity during task execution.

Neuroimaging evidence indicates that delivering pain during cognitive task execution does not produce a simple additive activation pattern but generates a distinct brain state (Bantick et al., 2002; Drevets & Raichle, 1998). For example, painful stimulation during Stroop task execution reduces activity in somatosensory cortex, SMA, MCC, insula, and cerebellum, while increasing subgenual ACC and OFC activation (Bantick et al., 2002; Seminowicz & Davis, 2007). The subgenual ACC and OFC are implicated in cognitive pain modulation, including mindful meditation (Zeidan et al., 2015). The aMCC and insula are key nodes of the salience network (Legrain et al., 2009; Wiech et al., 2010), whose activity is attenuated by reduced pain salience or hypnosis aimed at decreasing pain unpleasantness (Mouraux et al., 2011; Rainville et al., 1997). Although our design did not include ratings of a pain-only condition and therefore cannot directly assess task-driven pain modulation, our whole-brain results align with this literature. We observed increased subgenual ACC and OFC activation, decreased SMA, right somatosensory cortex and cerebellum activation, and deactivation in anterior insula and MCC. Increased activation in paracentral lobule, angular gyrus, precuneus, PCC, and lateral prefrontal regions likely reflects enhanced sensorimotor processing and cognitive control required to maintain performance under pain (Kimura et al., 2024; Schmidt et al., 2012).

Across parametric regression analyses, aMCC consistently showed negative associations with perceived effort. This pattern was observed independently of condition, in force-related contrasts (within- and between-participants), and in pain-related within-participants analyses. Notably, aMCC/MCC activity was never positively correlated with effort ratings. These findings contrast with accounts proposing that increased effort should involve enhanced cingulate activation (Mangin & Pageaux, 2026; Shenhav et al., 2016). Instead, reduced aMCC activity may reflect decreased motivational drive or subjective value associated with the ongoing action. Given the role of aMCC in cost evaluation and motivated behavior (Klein-Flügge et al., 2016; Le Heron et al., 2018), its reduced activity during higher perceived effort suggests that effort may be experienced as a costly state associated with diminished motivational value.

Importantly, the neural correlates of perceived effort differed when effort was assessed during high vs. low force production or under pain. During high-force contractions, perceived effort was associated with increased activity in SMA, primary motor cortex, and cerebellum, alongside decreased activity in aMCC and valuation-related regions. This pattern suggests that when greater force is required, perceived effort primarily reflects increased motor command and sensorimotor processing (de Morree et al., 2012). By contrast, under painful stimulation, higher perceived effort was associated with reduced activity in SMA, aMCC, and anterior insula, and increased activity in subgenual ACC, OFC, and frontoparietal regions. This pattern is consistent with engagement of pain modulation and cognitive control mechanisms (Bantick et al., 2002). The opposite direction of SMA modulation across force and pain contexts indicates that effort under pain cannot be reduced to amplified motor output. Instead, it likely reflects additional regulatory processes required to suppress pain salience and maintain task engagement.

Clinical studies identified the SMA as critical for initiating voluntary movement (Laplane et al., 1977), although the sense of volition remained intact after SMA lesion (Sjöberg, 2021). In contrast, lesions affecting the aMCC and striatum are associated with apathy, characterized by reduced motivation and goal-directed behavior (Le Heron et al., 2018), underscoring their role in motivational drive. In the present study, participants exerted effort without explicit reward. In this context, aMCC activity may reflect the low motivational value of the imposed task, whereas SMA activity may index the increased motor command necessary to perform high-force contractions (de Morree et al., 2012; Zénon et al., 2015). Perceived effort may therefore emerge from the combination of heightened motor command and reduced motivational drive. Although our design cannot disentangle these components, Croxson et al. (2009) provides a useful interpretive framework. In that study, SMA activity increased both during cues anticipating high task demand and cognitive task execution, independently of reward magnitude. By contrast, aMCC activity increased during cues predicting high reward, particularly with low task demand; but remained below baseline during task execution. This pattern supports the view that SMA may primarily encode effort magnitude, whereas aMCC may represent effort’s motivational value.

During task execution under pain, the functional contribution of aMCC and SMA may shift, given their central role in pain processing. The aMCC is a hub for the affective and motivational dimension of pain (Bushnell et al., 2013), as well as for sensory-integrative processes guiding pain avoidance behavior (Vogt et al., 1993) through connections with SMA (Hoffstaedter et al., 2014). However, when cognitive resources are allocated to task demands, pain salience may be attenuated and avoidance responses become less relevant. Under these conditions, reduced aMCC–SMA engagement circuitry may reflect downregulation of pain-related motivational and motor preparatory processes in favor of goal-directed performance (Shackman et al., 2011).

Whole-brain analyses independent of experimental condition revealed a complementary pattern: perceived effort positively covaried with activity in sensorimotor and visuomotor integration regions (post-central gyrus, precuneus, cerebellum), while negatively covarying with vmPFC, ventral striatum, PCC, and OFC. The vmPFC and PCC are central nodes of default mode networks (Wang et al., 2020), whose stronger deactivation during higher effort may reflect increased externally directed task engagement (Weissman et al., 2006). Given the role of vmPFC–striatal circuitry in subjective value encoding (Padoa-Schioppa, 2011), reduced activity during higher perceived effort may indicate that effort reflects a state where the cost of performing the task increases relative to perceived benefits. In such a framework, effort perception may emerge from a dynamic trade-off between intensifying motor command signals and diminishing value signals, consistent with theoretical accounts positioning effort as a cost variable guiding adaptive behavior (Shenhav et al., 2017).

Taken together, our findings indicate that maintaining task performance during pain is effortful and engages cost-benefit processes. While task performance remained stable, perceived effort increased, and this subjective cost was associated with systematic brain activity modulation. Across analyses, aMCC and SMA emerged as key, context-dependent regions. Neural patterns differed depending on whether perceived effort was altered by increased force level or nociceptive interference, underscoring that effort perception is influenced by cognitive and motor processes (Villeneuve et al., 2025). Although our design does not allow causal inferences or direct assessment of pain-only modulation, associations between perceived effort and brain activation patterns suggest that effort perception may reflect monitoring of motor and cognitive resources engaged in task execution, together with changes in the motivational value assigned to the action. Reduced activity in valuation-related regions alongside context-specific modulation of motor regions further supports the view that effort integrates both cost-related and motivational components. In contexts where actions are not explicitly rewarded, increased perceived effort may function as a negative reinforcement signal, biasing future behavior away from demanding, non-rewarding activities (Akaishi et al., 2016). Future research should determine how these mechanisms unfold over longer timescales and whether similar neural signatures characterize clinical conditions with altered effort and marked by chronic pain, fatigue, or motivational disturbances (Jurgelis et al., 2021; Salamone et al., 2016; Van Damme et al., 2018).

## Author Contributions

IM: conceptualization, methodology, software, validation, formal analysis, investigation, data curation, writing – original draft, writing – review & editing, visualization, project administration. MEP: methodology, validation, formal analysis, data curation, visualization, writing – original draft, writing – review & editing. TM: conceptualization, methodology, software, investigation, writing – review & editing. MB: formal analysis, investigation, visualization, writing – review & editing. MG: conceptualization, writing – review & editing. SB: conceptualization, writing – review & editing. RO: conceptualization, writing – review & editing, funding acquisition. JIC: formal analysis, investigation, data curation, resources, project administration, writing - review & editing. MR: conceptualization, writing - review & editing, funding acquisition. PR: conceptualization, methodology, validation, resources, writing - review & editing, visualization, supervision, funding acquisition. BP: conceptualization, methodology, validation, resources, writing - review & editing, visualization, supervision, funding acquisition.

## Preprint servers

The manuscript was deposited as a preprint on the website bioRxiv at the link https://doi.org/10.64898/2026.04.17.719211.

## Data availability statement

All data files are available at https://zenodo.org/records/20445958.

## Acknowledgments

Funding

IM was supported by Bourse d’études du Réseau québécois de bio-imagerie, Bourse de Mérite aux Cycles Supérieurs de l’Université de Montréal and Bourse Fonds de recherche du Québec – Nature et technologies. MEP was supported by Bourse Fonds de recherche du Québec – Nature et technologies. TM was supported by the postdoctoral scholarship of the Centre de Recherche de l’Institut Universitaire de Gériatrie de Montréal (CRIUGM), and the postdoctoral research scholarship from the Fonds de Recherche du Québec – Nature et technologie. MB was supported by the Natural Sciences and Engineering Research Council of Canada through a doctoral postgraduate scholarship and the “Formation de doctorat” scholarship from the Fonds de recherche du Québec – Nature et technologie. BP research is supported by the Natural Sciences and Engineering Research Council of Canada – Discovery Grant and the Chercheur Boursier Junior 1 from the Fonds de recherche du Québec – Santé. This project was also funded by a Pilot Project Grant from the Réseau Québecois de Recherche sur la Douleur, and was part of an international collaboration between BP and SB supported by the XIe Commission mixte permanente Québec–Wallonie-Bruxelles.

## Additional Contributions

We would like to thank Andre Cyr for technical support and Julie Boyle for assistance with the open-access process.

## Use of AI

The authors used ChatGPT to assist with English language editing and grammar improvement during manuscript preparation. After using this tool, the authors reviewed and edited the content as needed and take full responsibility for the content of the published article. All scientific content, data interpretation, and conclusions were generated solely by the authors.

